# The Clinical Significance and Biological Function of *DPEP1* in B-cell Acute Lymphoblastic Leukemia

**DOI:** 10.1101/742551

**Authors:** Jia-Min Zhang, Yan Xu, Robert Peter Gale, Li-Xin Wu, Jing Zhang, Yong-Huai Feng, Ya-Zhen Qin, Hao Jiang, Qian Jiang, Bin Jiang, Yan-Rong Liu, Yu-Hong Chen, Yu Wang, Xiao-Hui Zhang, Lan-Ping Xu, Xiao-Jun Huang, Kai-Yan Liu, Guo-Rui Ruan

## Abstract

Dehydropeptidase-1 (*DPEP1*) is a zinc-dependent metalloproteinase abnormally expressed in many cancers. However, its potential role in adults with B-cell acute lymphoblastic leukaemia (ALL) is unknown.

We found that in adults with common-B-cell ALL high *DPEP1* transcript levels at diagnosis was independently-associated with an increased CIR and worse RFS compared with subjects with low transcript levels. We show an increased proliferation and pro-survival role of *DPEP1* in B-cell ALL cells *via* regulation of phosphCREB and p53 which may be the biological basis of the clinical correlation we report. Our data implicate *DPEP1* expression in the biology of common B-cell ALL in adults. We report clinical correlates and provide a potential biological basis for these correlations. If confirmed, analyzing *DPEP1* transcript levels at diagnosis could help predict therapy-outcomes. Moreover, regulation of *DPEP1* expression could be a therapy target in B-cell ALL.

## 1. Introduction

Relapse is the leading cause of therapy-failure in adults with B-cell ALL. Many pre-therapy variables correlate with relapse risk but these are inaccurate in many instances and explain only about one-half of the variance in relapse outcomes. Consequently, new variables independently-correlated with relapse risk are needed. Dehydropeptidase-1 (*DPEP1*) is a zinc-dependent metalloproteinase. *DPEP1* deletions are common in several cancers^1^. *DPEP1* activity regulates concentrations of glutathione, leukotrienes and pro-inflammatory molecules associated with cancer development^2^. Several studies assessed *DPEP1* transcript levels in diverse cancers. In some, transcript levels were prognostic and/or predictive of outcomes^3–5^. However, the role and clinical relevance of *DPEP1* expression in cancer is controversial and molecular mechanism by which it acts may vary between cancers^4, 6–8^.

Persons with B-cell ALL have high *DPEP1* transcript levels compared with normals^9^. However, the functional implication and clinical relevance of this observation, if any, is unknown. We evaluated *DPEP1* transcript levels in bone marrow samples from adults with newly-diagnosed B-cell ALL. We found high *DPEP1* transcript levels were associated with worse clinical outcomes. We also studied *DPEP1* expression in a human B-cell ALL cell line and in a mouse xeno-transplant model. Our data suggest *DPEP1* transcript levels independently-correlate with a higher cumulative incidence of relapse CIR and worse relapse-free survival (RFS) in adults with common B-cell ALL.

## 2. Materials and Methods

### 2.1. Subjects

Bone marrow samples were obtained from 235 consecutive adults (>16 years) with new-diagnosed, untreated B-cell ALL and 28 normals at the Hematology Department of Peking University Peoples Hospital March, 2009 to March, 2016. Subjects were followed until death, loss to follow-up or June, 2018. Risk-stratification and therapy details are published^10, 11^.

Subjects with *BCRABL1* (N=91) also received tyrosine kinase-inhibitors^11^. Subjects achieving complete remission (N=208; 85%) were included in analyses of predictive variables. Because prognosis depends on ALL subtype, we pre-specified a separate analysis of CIR and RFS in subjects with common B-cell ALL (N=167). There were too few subjects with pre-B-cell ALL (N=14) and pro-B-cell ALL (N=27) to separately analyze for CIR and RFS. 105 subjects with common-B-cell ALL received an allotransplant 21 from an HLA-identical sibling and 84, from a HLA-haplotype-matched relative.^12, 13^. The study was approved by the Ethics Committee of Peking University Peoples Hospital. Written informed consent was obtained according to the Declaration of Helsinki.

### 2.2. Leukemia cell lines

Human leukemia cell lines, including SUP-B15, BALL-1, Nalm6, KG-1, NB4, HL60, MEG-01, K562, MOLP2, SKO-007, U937, Raji, Ramos and IM-9, were purchased from American Type Culture Collection (ATCC, Manassas, VA, US) and cultured in RPMI-1640 medium containing 10% heat-inactivated fetal calf serum, penicillin and streptomycin (all from Sigma-Aldrich. St. Louis, MO, US) at 37°C with 5% CO_2_ in a humidified incubator. An EBV-transformed normal human B-cell line was a gift from Dr. Jiang-Ying Liu of Beijing Key Laboratory of Haematopoietic Stem Cell Transplantation.

### 2.3. Immune phenotype, measurable residual disease (MRD) and cytogenetic analyses

Bone marrow samples were analyzed using standard 4-color flow cytometry (FCM)^14^. 3 immune phenotypes were identified:(1) early precursor B-cell ALL (pro-B-ALL): CD10-negative, CD19-positive, cCD79a-positive, cCD22-positive and TdT-positive; (2) common B-cell ALL: CD10-positive; and (3) precursor B-cell ALL (pre-B-ALL): cytoplasmic μ-positive, sIg-negative, CD10-positive or negative^11^. MRD was quantified by analyzing leukemia-associated aberrant immune phenotypes (LAIPs) by4-color flow cytometry as described^15^. A positive MRD-test was defined as ≥0.1% of cells with an LAIP phenotype in ≥1 bone marrow sample in 1^st^ complete remission^15^. Cytogenetic analyses were performed by G-banding^16^. *BCRABL1* transcripts and *KMT2A (MLL)* rearrangements were detected with quantitative real-time polymerase chain reaction (RT-qPCR) ^17, 18^. *IKZF1* deletion was detected as described^10^. Risk cohort was defined as previously described^11^.

### 2.4. Lentiviral transduction

BV173 or BALL-1 cells were infected with human *DPEP1* over-expression lentivirus or *DPEP1* shRNA lentivirus or corresponding blank control (GeneChem, Shanghai, China, MOI=100). Media containing lentiviral particles were replaced with complete medium 12 h post-infection and stably-transfected BV173 cells were selected with 0.5 μg/ml puromycin dihydrochloride (GeneChem) for ≥6 w post-infection. *DPEP1* expression levels were confirmed by RT-qPCR and western blotting.

### 2.5. RNA preparation and RT-qPCR

Mononuclear cells were isolated from bone marrow samples by Ficoll-Hypaque™ density gradient centrifugation. RNA extraction and cDNA synthesis were performed as described. mRNA expression levels were determined by TaqMan^19 ®^. *DPEP1* transcript levels were normalized to *ABL1* expression as recommended^20^. Copy numbers of *DPEP1* and *ABL1* were calculated from standard curves using Ct values. Samples were assayed in triplicate and average threshold Ct values calculated.

### 2.6. Western blotting

Cells were lysed in the presence of protease cocktail inhibitor, soluble lysates electrophoresis on SDS-polyacrylamide gels and transferred to a polyvinylidene fluoride membrane (Bio-Rad, Hercules, CA, US). Membranes were probed with specific antibodies and signals visualized using Odyssey Infrared Imaging System (LI-COR Biosciences, Lincoln, NE, US). GAPDH was used as a loading control.

### 2.7. Cell proliferation and viability assay

Proliferation was determined with the Cell Counting Kit-8 assay (CCK8, Dojin Laboratories, Kumamoto, Japan). Briefly, cells were seeded into 96-well plates at a density of 4×10E4 according to the manufacturer’s instructions. Plates were scanned by a microplate reader at 450 nm at indicated time points. CCK8 was also used to determine cell viability after exposure to drugs, including daunorubicin, dexamethasone, methotrexate, cytarabine and imatinib (Solarbio, Beijing, China). Experiments were performed in triplicate.

### 2.8. Cell-cycle analyses

The cells were seeded onto 6-well plates and starved by adding serum-free medium for G_1_ synchronization. After 24 h medium containing 10% fetal bovine serum was added for 48 h. Cells were fixed in 75% ethanol, stained with propidium iodide (BD Pharmingen, San Jose, CA, US) and analyzed by flow cytometry. Results were analyzed with ModFit LT2.0 software (Coulter Electronics, Hialeah, FL, USA).

### 2.9. Detection of apoptosis

Apoptosis was determined using the Annexin V-PI or Annexin V-7AAD Apoptosis Detection Kit (Sigma-Aldrich) according to the manufacturer’s instructions. Samples were immediately analyzed by flow cytometry (Beckman Coulter). Approximately 5×10E+5 cells were analyzed in each sample.

### 2.10. Colony-forming assays

Cells were suspended in 1 mL of complete MethoCult™ medium and plated in 6 well plates at a concentration of 4 × 10E+3 cells/well. Colonies were maintained at 37°C with 5% CO_2_ and 95% humidity for 7d, counted and scored after staining with 1% crystal violet (Sigma). Colonies of ≥50 cells were scored. Assays were performed in triplicate.

### 2.11. Cell migration and invasion assay

Cells were seeded into the upper chamber of a trans-well insert (pore size, 8 μm, Corning, Corning, NY, US) in RPMI-1640 supplemented with 1% FBS. The upper chamber was placed into the trans-well containing medium supplemented with 10% FBS in the lower chamber. For the invasion assay, a Matrigel coating (BD systems, San Jose, CA, US) was used. Cells remaining in the lower surface of the insert filter 24 h were stained with crystal violet. Experiments were done in triplicate.

### 2.12. Tumor xenograft mouse model

Male athymic 6-week-old Balb/c nude mice (Beijing HFK Bioscience Co., Ltd.; Beijing, China) were pretreated with intraperitoneal injections of cyclophosphamide once daily at 100 mg/kg for 2 consecutive d. BV173 cells (5×10E+6 cells in 0.1 mL of PBS) transduced with the indicated lentivirus were injected subcutaneously into the dorsal right flank area (5 mice/group). Tumor diameters were measured every 2 d for 15 d. Tumor volume (mmE+3) was estimated by measuring the longest and shortest diameters of the tumor as described [49]. Mice were euthanized on day 15 and tumors removed for subsequent experiments. Animal experiments were approved by the Animal Ethics Committee of Peking University Health Science Center.

### 2.13. Statistical analyses

Complete remission, relapse and refractory disease were defined as described^11^. Cumulative incidence of relapse (CIR) was calculated from date of 1^st^ complete remission to 1^st^ relapse or death in complete remission. Relapse-free survival (RFS) was determined from dates of 1^st^ complete remission to 1^st^ relapse. Subjects achieving complete remission (N=208) were included in analyses of predictive variables for CIR and RFS. Differences across groups were compared using Pearson Chi-square analysis or Fisher exact test for categorical data and the Mann-Whitney U test or Student t-test for continuous variables. Receiver operating characteristic (ROC) curves were constructed to evaluate sensitivity and specificity of *DPEP1* transcript levels to estimate CIR. Youden Index was used to calculate the optimal cutoff for high and low transcript levels^21^. Survival functions were estimated by the Kaplan-Meier method and compared by the log-rank test. A Cox proportional hazard regression model was used to determine correlations between *DPEP1* transcript levels, CIR and RFS. Variables with a *P*-value <0.10 in univariable analyses were entered into the multivariable analyses including age (≥ vs. <35 years), sex (M/F), WBC at diagnosis (<vs. ≥30×10E+9/L), *BCRABL1* (Negative/Positive), *MLL* translocation (N/Y), *IKZF1* deletion (N/Y), transplant (N/Y), MRD-test result in complete remission (Negative/Positive) and *DPEP1* transcript level at diagnosis (High/Low). A *P*<0.05 was considered significant in multivariable analyses. The sensitivity analysis was performed with only subjects who received the appropriate designated therapy (high-risk or standard risk with positive MRD: transplant; standard risk with negative MRD: no transplant;) included. Analyses were performed by SPSS software version 18.0 (Chicago, IL, USA), GraphPadPrism™ 5.01 (San Diego, California, USA) and R software package (version 3.1.2; http://www.r-project.org). Median follow-up is 23 months (range, 1–104 months).

## 3. Results

### 3.1. *DPEP1 is highly-expressed in bone marrow lymphoblasts* from subjects *with B-cell ALL and correlates with disease state*

Transcript levels of *DPEP1* in bone marrow samples were assessed in 235 consecutive adults with newly-diagnosed, untreated B-cell ALL, 28 normals and a normal human B-cell line. Transcript levels were higher in the B-cell ALL subjects compared with normals and with a normal-B cell line (Figure 1S-A). We then compared *DPEP1* transcript levels in subjects with B-cell ALL in complete remission and relapse. Transcript levels decreased to a level like normals in subjects in complete remission but were increased at relapse (Figure 1S-A). Sequential determinations of *DPEP1* transcript levels in 9 subjects are displayed in Figure 1S-B.

### 3.2. DPEP1 transcript level is independently associated with CIR and RFS in subjects with common B-cell ALL

To interrogate the predictive impact of *DPEP1* transcript levels on outcomes, we analyzed CIR and RFS in subjects with common B-cell ALL achieving complete remission. Characteristics were summarized in Table 1S. High *DPEP1* transcript levels at diagnosis were independently-associated with increased CIR (HR=2.05 [1.15,3.63]; *P*=0.014) and worse RFS (HR=1.96 [0.10, 0.36]; *P*=0.013; Table 1). Other variables significantly correlated with CIR and RFS were a positive MRD-test in complete remission (HR=2.23 [1.27, 4.00]; *P*=0.005; and HR=2.15 [1.25, 3.58]; *P*=0.005), receiving a transplant (HR=0.19 [0.10, 0.36]; *P*=0.000; and HR=0.22 [0.12, 0.39]; *P*=0.000) and having *BCRABL1* (HR=1.65 [0.93, 2.94]; *P*=0.085; and HR=1.95 [1.14, 3.36]; *P*=0.016, Table 2).

**Table 1.** Multivariable analyses of CIR and RFS in adults with common B-cell ALL

|  | HR (95% CI) | *P*-value |
| --- | --- | --- |
| CIR |  |  |
| *DPEP1* transcripts (High/Low) | 2.05 (1.15, 3.63) | 0.014 |
| Transplant (Y/N) | 0.19 (0.10,0.36) | <0.0001 |
| MRD-test (+/-) | 2.23 (1.27,4.00) | 0.005 |
| *BCRABL1*(+/-) | 1.65 (0.93,2.94) | 0.085 |
| RFS |  |  |
| *DPEP1* transcripts (High/Low) | 1.96 (1.15,3.33) | 0.013 |
| Transplant (Y/N) | 0.22 (0.12,0.39) | <0.0001 |
| MRD-test (+/-) | 2.15 (1.25,3.58) | 0.005 |
| *BCRABL1*(+/-) | 1.95 (1.14,3.36) | 0.016 |

Abbreviations: HR, hazard ratio; CI, confidence interval; MRD, measurable residual disease.

Our analyses were confounded by protocol violations. To address this bias, we conducted sensitivity analyses. Only *DPEP1* transcript level remained significant in sensitivity analyses when only subjects who received the appropriate designated therapy (high-risk or standard risk with positive MRD: transplant; standard risk with negative MRD: no transplant) were included (CIR: HR=2.63 [1.10, 6.25]; *P*=0.029; and RFS: HR=2.48 [1.15, 5.36]; *P*=0.021).

### 3.3. DPEP1 promotes the proliferation and survival of leukemia cell lines

To determine the functional significance of *DPEP1* expression *in vitro* we used a lentivirus vector to knock down *DPEP1* in the BV173 cell line and *DPEP1* over-expressing (OE) cell lines were derived from BV173 and BALL-1 cells. The over-expression and knockdown efficiency were verified by RT-PCR (Figure 1A) and western blot analyses (Figure 1B).

**Figure 1.**
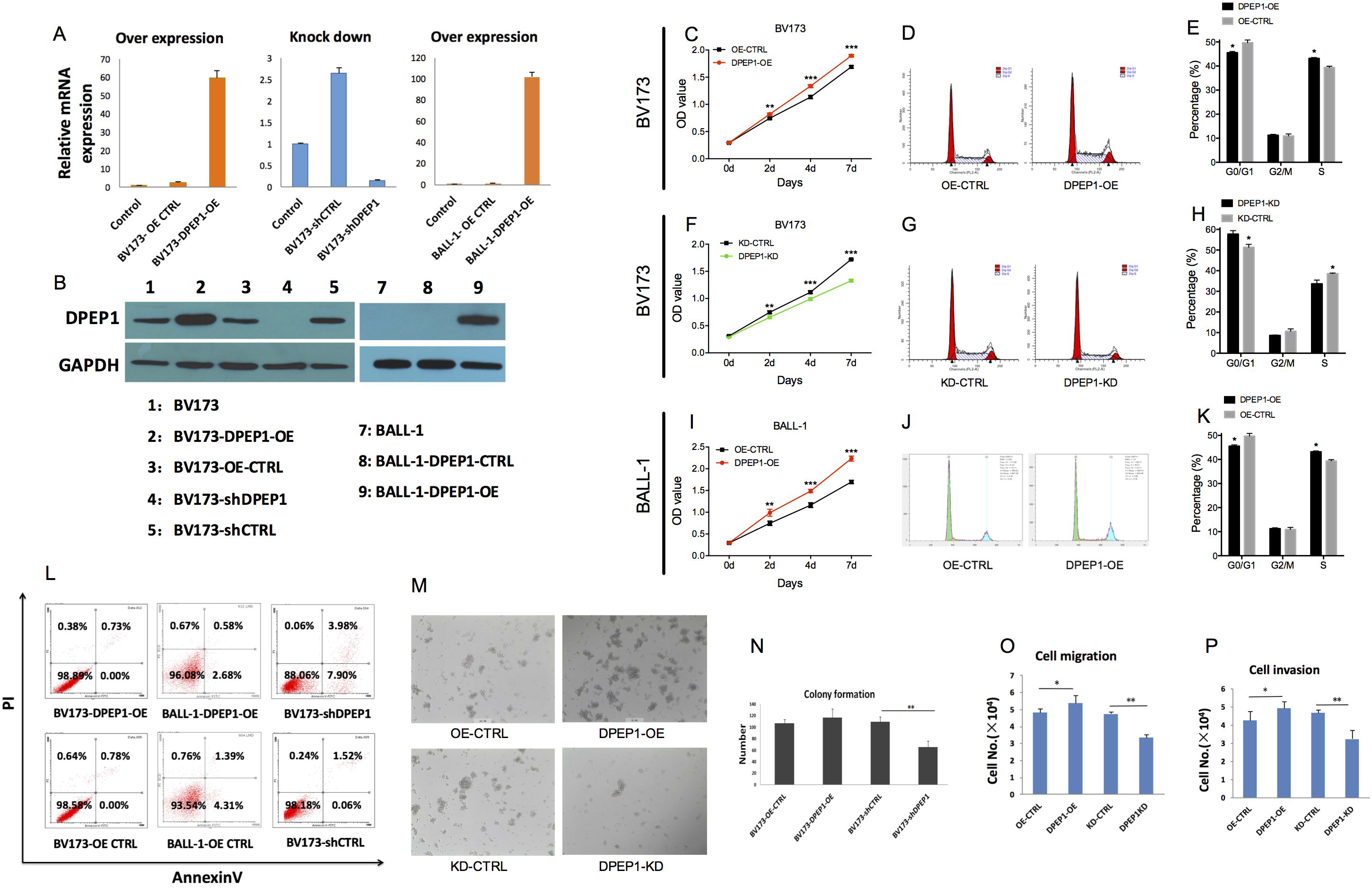
*DPEP1* transcription promotes proliferation and survival of leukemia cells. Cell proliferation detected by CCK-8 in BV173 PEP1-OE cells (A), BV173 *DPEP1*-KD cells (D) and BALL-1 *DPEP1*-OE cells (G) with corresponding vector controls. Cell cycle analysis in BV173 *DPEP1*-OE cells (B, C), BV173 *DPEP1*-KD cells (E, F) and BALL-1 *DPEP1*-OE cells (H, I) with their respective vector controls. (J) the apoptosis of *DPEP1*-OE and *DPEP1*-KD BV173 cells. Colony formation in *DPEP1*-OE and *DPEP1*-KD BV173 cells. Cell migration (M) and cell invasion (N) detected by transwell assays using *DPEP1*-modified BV173 cells. *, *P*< .05; **, *P*< .01; ***, *P*< .001. Error bars indicate the SD. *DPEP1*, dehydropeptidase1; CTRL, control; OE, over-expression; KD, knockdown.

We first determined whether *DPEP1* could alter proliferation and cell-cycle of leukaemia cells. BV173 cells over expressing *DPEP1* displayed increased proliferation (Figure 1C) and more rapid cell-cycle progression (Figure 1D, E) compared with controls. Knockdown of *DPEP1* resulted in a significant decrease in proliferation (Figure 1F) and slowed cell-cycle progression in BV173 cell lines (Figure 1G, H). The effect of *DPEP1* was further confirmed in *DPEP1*-OE BALL-1 cell lines (Figure 1I, J and K). Next, we detected whether *DPEP1* could influence cell death *in vitro*. Apoptosis was relatively rare in control cell lines and *DPEP1*-OE did not markedly affect this rate (Figure 1L). However, *DPEP1*-KD significantly increased apoptosis in the BV173 cell line(Figure 1L). To confirm the role of *DPEP1* in the survival and proliferation of B-cell ALL cells we performed clonality assays. Colony numbers were markedly increased in *DPEP1*-OE cells but decreased in *DPEP1*-knockdown cells compared with vector controls (Figure 1M, N). In addition, migration (Figure 1O) and invasion (Figure 1P) were significantly enhanced in *DPEP1*-OE BV173 cells but attenuated in *DPEP1*-knockdown cells detected by transwell assays. These data suggest *DPEP1* transcription favors proliferation and survival of B-cell ALL cell lines.

### 3.4. DPEP1 improves the survival of leukaemia cells exposed to anti-leukaemia drugs

We next studied interactions between *DPEP1* expression and cell viability in cultures with anti-leukaemia drugs including dexamethasone, daunorubicin, cytarabine and imatinib (BV173 is a *BCRABL1*-positive). *DPEP1*-OE B-ALL-1 cells were more viable after cultures with dexamethasone, daunorubicin or imatinib compared with controls (Figure 2A-D). *DPEP1*-OE BV173 cells showed the same trend of increased viability in the presence of anti-leukaemia drugs (Figure 2E-H), whereas *DPEP1*-KD BV173 cells had a significant decrease in viability under these conditions (Figure 2I-L).

**Figure 2.**
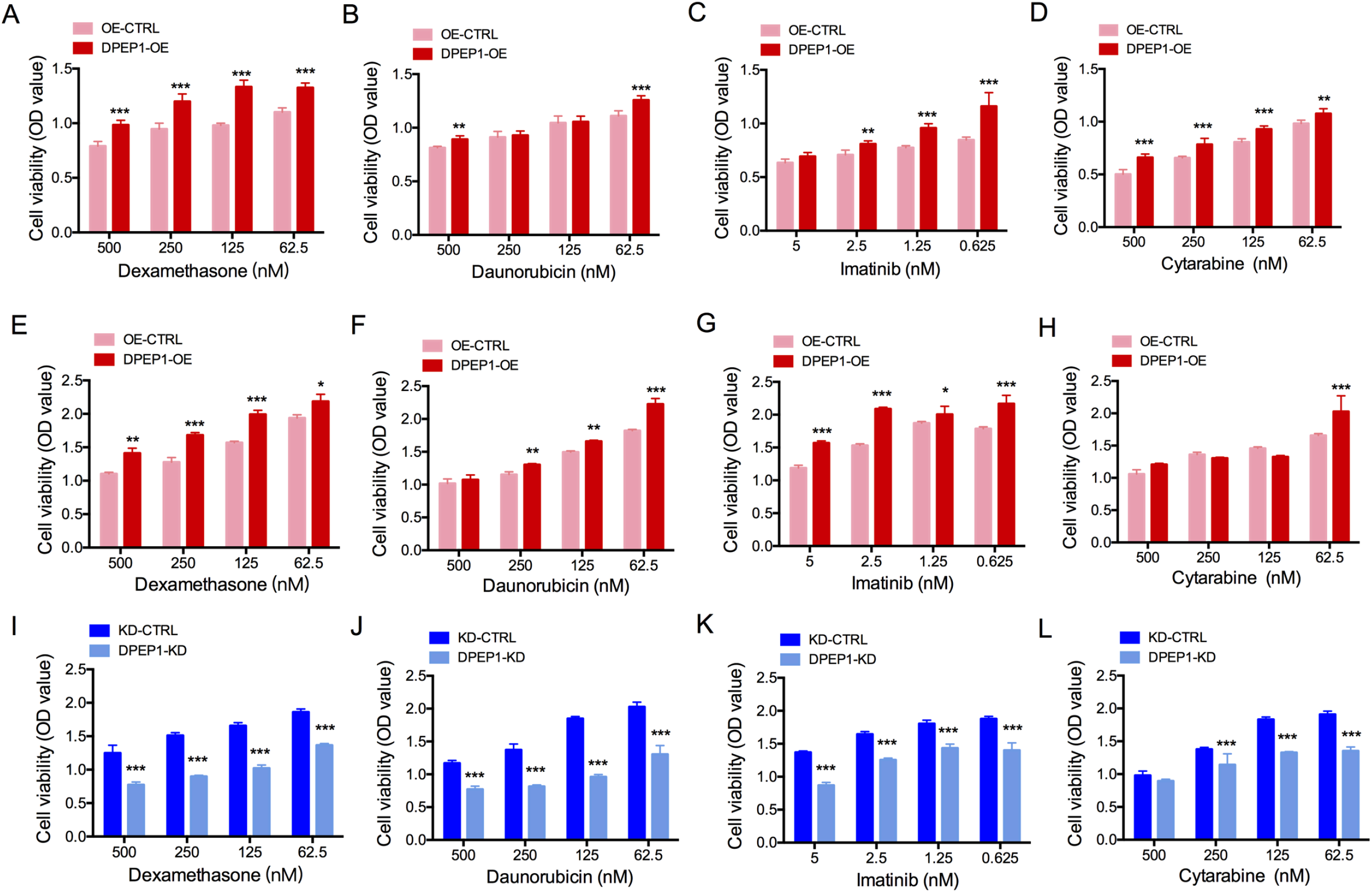
*DPEP1* over-expressionand anti-leukaemia drug exposure. *DPEP1* over-expression increased proliferation of BALL-1 cells exposed to dexamethasone (A), daunorubicin (B), imatinib (C) and cytarabine (D) compared with controls. *DPEP1* over-expression increases survival of BV173 cells exposed to dexamethasone (E), daunorubicin (F), imatinib (G) and cytarabine (H) compared with controls. *DPEP1* knockdown decreased survival of BV173 cells to dexamethasone (I), daunorubicin (J), imatinib (K) and cytarabine (L) compared with controls. *, *P*< .05; **, *P*< .01; ***, *P*< .001. Error bars indicate the SD. *DPEP1*, dehydropeptidase1; CTRL, control; OE, over-expression; KD, knockdown.

### 3.5. DPEP1 promotes the growth and survival of leukemia cells in vivo

Xeno-transplants were done with *DPEP1*-OE and *DPEP1*-KD cells to further investigate *in vivo* effects of *DPEP1* expression. Knockdown of *DPEP1* markedly decreased tumor growth compared with controls (Figures 3A and B). In contrast, tumors were larger in *DPEP1*-OE cells (Figure 3A and B) compared with controls. Histological examination showed more abundant blood sinuses in *DPEP1*-OE cell-derived tumors but a larger necrotic area in tumors derived from *DPEP1*-KD cells (Figure 3C). TUNEL staining confirmed increased apoptosis in specimens from the *DPEP1*-KD group compared with those from the vector controls corroborating the *in vitro* findings on apoptosis (Figure 3D).

**Figure 3.**
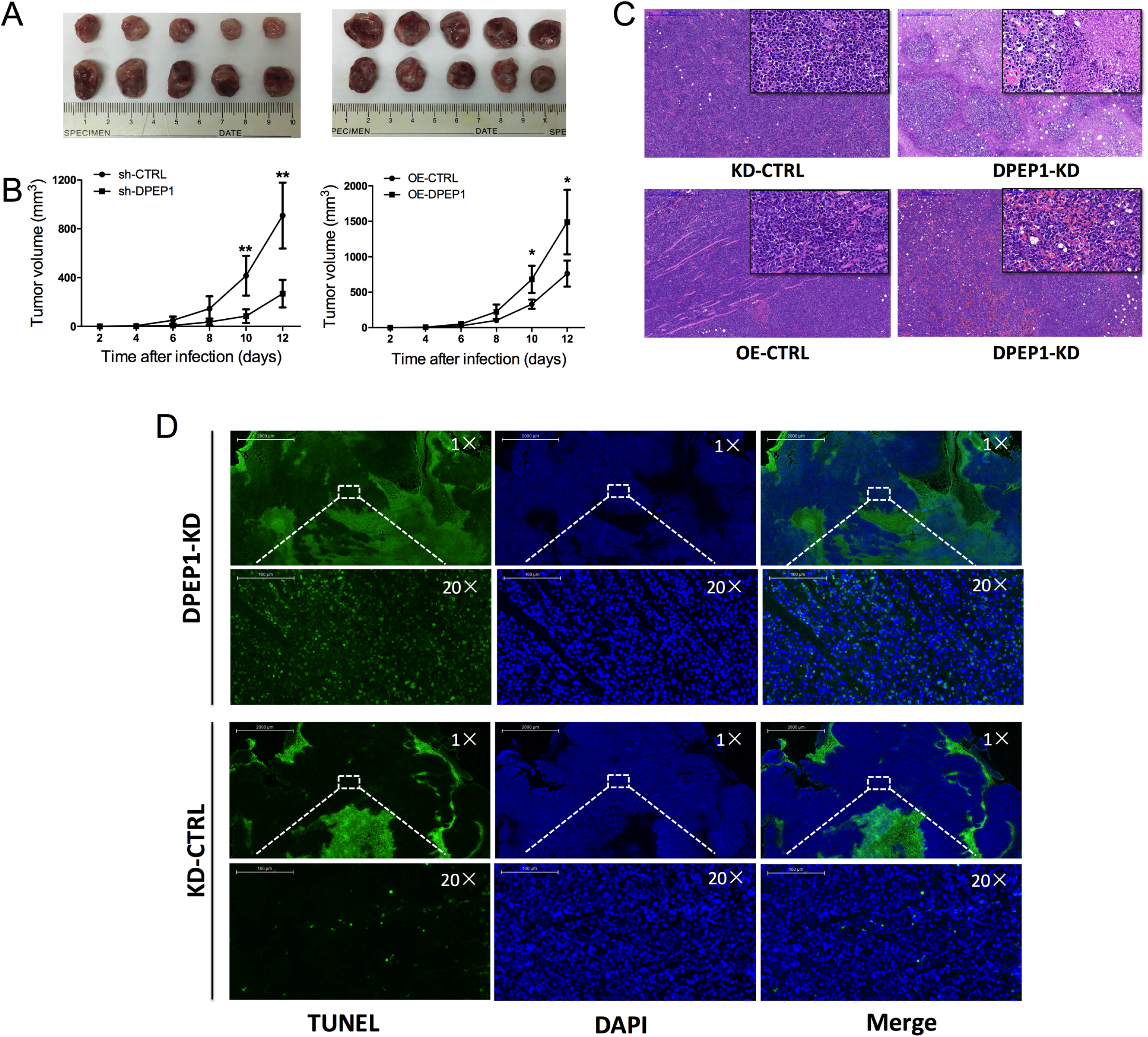
*DPEP1* expression and growth and survival of leukaemia cells in a xeno-transplant mouse model. (A-B) *DPEP1*-OE promotes tumor growth whereas *DPEP1*-KD decreased tumor growth; (B) *DPEP1*-OE tumors had a high density of cells with abundant blood sinuses whereas *DPEP1*-KD tumors had more necrosis in HE sections (magnification 5X, scale bar=500um; magnification 40X, scale bar= 50um). (D) TUNEL staining showed a stronger trend of apoptosis in *DPEP1*-OE tumors compared with controls (64.5± 10.3 vs. 16.7 ± 3.4; *P*< 0.001, magnification 1X, scale bar= 2000μm; magnification 20X, scale bar= 100μm). *, *P*< .05; **, *P*< .01; ***, *P*< .001. Error bars indicate the SD. *DPEP1*, dehydropeptidase1; CTRL, control; OE, over-expression; KD, knockdown; TUNEL, terminal-deoxynucleotidyl transferase-mediated nick end labeling; DAPI, 4,6-diamino-2-phenyl indole.

### 3.6. Pathways involved in DPEP1-mediated alterations

To gain mechanistic insight into potential effector molecules involved in the increased proliferation and survival resulting from *DPEP1* expression, we examined changes in several leukemia cell growth- and apoptosis-related proteins in BV173 cells. Levels of p-mTOR and CDK4 were largely unaffected by *DPEP1* expression (Figure 4A). Phosphorylation of ERK was down-regulated in *DPEP1*-OE and *DPEP1*-KD cells and strongly inhibited in KD cells (Figure 4A). Phosphorylation of p38 was not markedly-affected in *DPEP1*-KD cells but was blocked in *DPEP1*-OE cells. Proliferation-related molecules p-AKT and p-CREB, which are activated through p-AKT, were up-regulated with *DPEP1* over-expression and blocked by *DPEP1* knockdown (Figure 4A). Level of pro-apoptotic p53 was increased after *DPEP1* inhibition although there were no changes in *DPEP1*-OE cells consistent with the apoptosis assessments *in vitro* and *in vivo* (Figure 4A).

**Figure 4.**
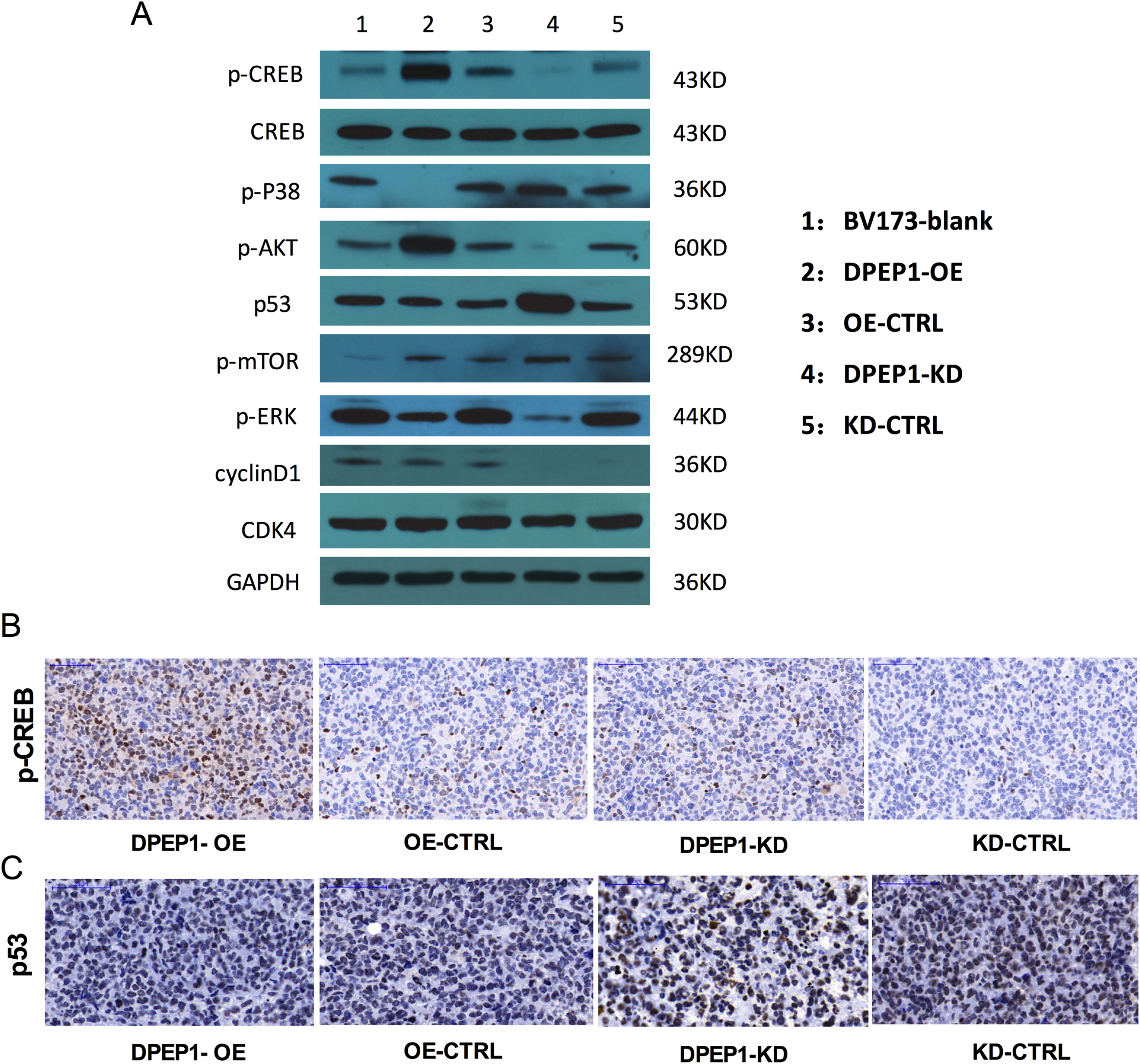
Changes in proliferation- and apoptosis-related proteins in *DPEP1* gene-modified BV173 cells and tumors. (A) Western blot analysis of proliferation- and apoptosis-related proteins in *DPEP1*-OE and *DPEP1*-KD BV173 cells and controls; (B) Immune histochemistry of pCREB and p53 in tumors derived from *DPEP1* gene-modified BV173 cells (magnification 20X, scale bar 50μm). *DPEP1*, dehydropeptidase1; CTRL, control; OE, over-expression; KD, knockdown.

Next, we studied p-CREB and p53 expression in mouse tumors. Immune histochemistry studies detected high levels of pCREB in *DPEP1*-OE tumors compared with controls (Figure 4B). However, p53 was highly expressed in *DPEP1*-KD tumors compared with tumors from the other groups (Figure 4C). These data suggest *DPEP1* silencing favors an *apoptotic, growth-inhibition* phenotype whereas *DPEP1*-OE cells favors a *proliferative* phenotype.

### 3.7. Activation of pCREB pathway is responsible for DPEP1-mediated proliferation and anti-apoptosis in B-cell-ALL cell lines

To confirm the role of pCREB in *DPEP1*-mediated effects, we inhibited pCREB in BV173 *DPEP1*-OE cells with KG-501 (Figure 5A). pCREB-inhibition (12.5μM) reversed the increased proliferation of *DPEP1*-OE cells (Figure 5B). This effect was confirmed in cell-cycle analyses where a decrease in the proportion of KG-501-treated *DPEP1*-OE cells in S-phase (Figure 5C). Conversely, blocking pCREB induced an apoptotic phenotype of BV173 cell lines (Figure 5D). These data suggest phosphorylation of CREB by increased expression of *DPEP1* is responsible for the changes in the growth and apoptosis of *DPEP1*-modified BV173 cell lines.

**Figure 5.**
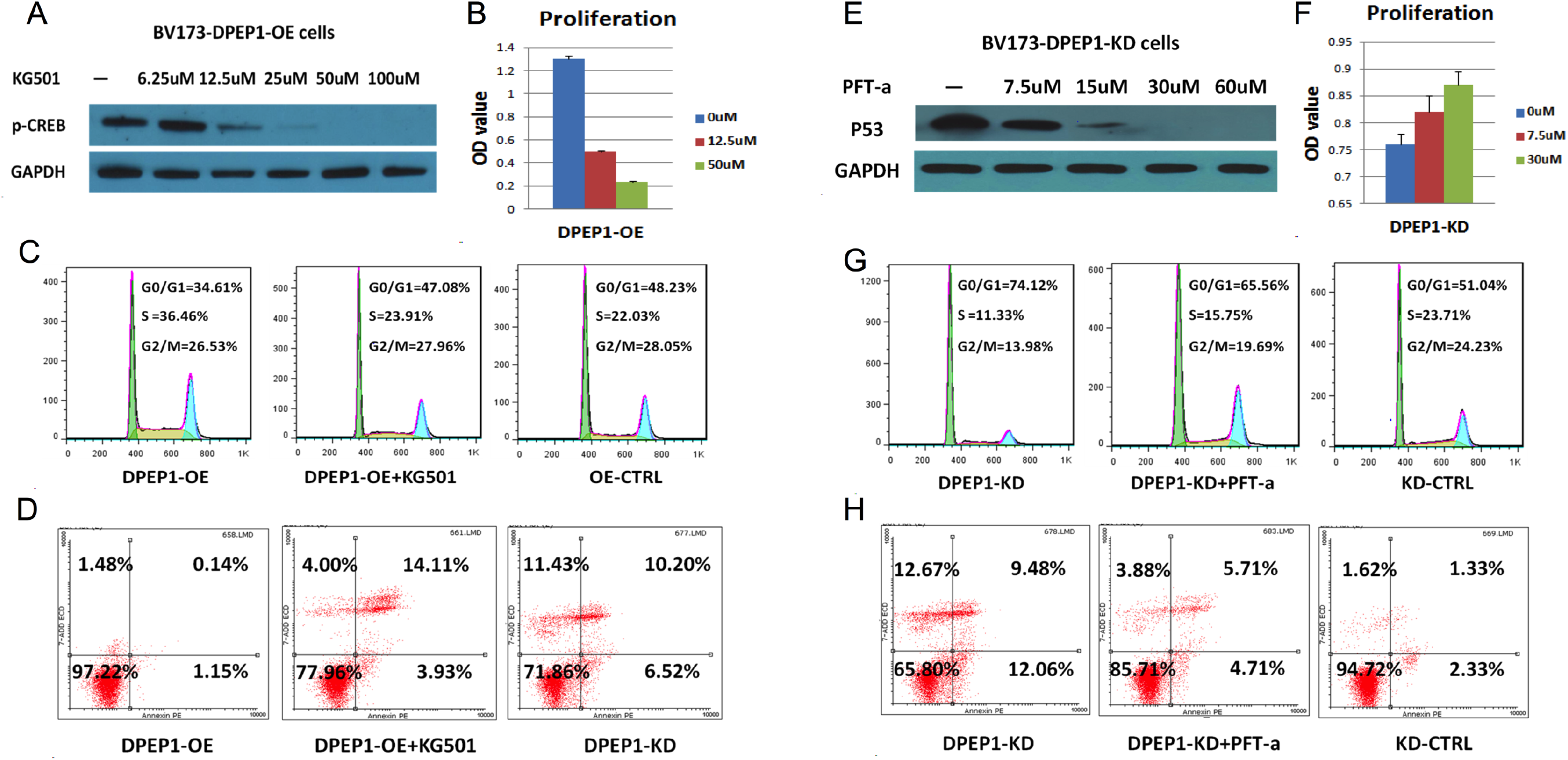
Inhibition of pCREB and p53 in *DPEP1*-modified BV173 cells. (A) Inhibition of p-CREB by KG-501 in *DPEP1*-OE BV173 cells. (B) Inhibition of pCREB and cell proliferation. (C) Inhibition of pCREB by KG-501 (12.5 μM) in *DPEP1*-OE cell and cycle progression; (D) Inhibition of pCREB by KG-501 (50 μM) and apoptosis. (E) Inhibition of p53 by PFT-a in *DPEP1*-KD BV173 cells; (F) Inhibition of p53 by PFT-a in BV173. (G) Inhibition of p53 by PFT-a (7.5 μM) and cell cycle; (H) Inhibition of p53 by PFT-a (7.5 μM) and apoptosis in *DPEP1*-KD BV173 cells. *P*< .05; **, *P*< .01; ***, *P*< .001. Error bars indicate the SD. *DPEP1*, dehydropeptidase1; CTRL, control; OE, over-expression; KD, knockdown; PFT-a, Pifithrin-α.

### 3.8. p53 is involved in DPEP1 silencing-induced apoptosis and growth inhibition

To verify the effect of p53 in *DPEP1*-KD cells we used PFT-a, a p53 inhibitor. Inhibition of p53 significantly reversed the proliferation defect in *DPEP1*-KD cells (Figure 5E). Treatment with PFT-a slightly reversed decreased proliferation in *DPEP1*-KD cells (Figure 5F). Similarly, suppression of p53 promoted progression of quiescent cells (G_0_/G_1_ phase) into S and G_2_ phase (Figure 5G). The pro-apoptotic phenotype observed in *DPEP1*-KD cells was markedly decreased by PFT-a treatment (Figure 5H). Taken together, these data indicated a critical role for p53 in *DPEP1* silencing-induced apoptosis and growth defects in BV173 cells.

## 4. Discussion

We found markedly-increased *DPEP1* transcript levels at diagnosis in bone marrow lymphoblasts from adults with B-cell ALL compared with normals. Transcript levels decreased in subjects achieving a complete remission but increased up on relapse. In multivariable analyses *DPEP1* transcript levels at diagnosis, a positive MRD-test result in complete remission, not receiving a transplant and having *BCRABL1* were significantly-associated with a higher CIR and worse RFS. Failure to detect significant correlations between predictive variables such as cytogenetics, WBC levels at diagnosis and *IZKF1*-deletion with CIR and/or survival is likely explained by 2 considerations: (1) much of the impact of these variables operates on the likelihood of achieving a complete remission whereas we analyzed only subjects in complete remission; (2) subjects with high-risk features such as adverse cytogenetics received different therapies than subjects without these features such as an transplant and/or tyrosine kinase-inhibitor therapy.

Our analyses of variables associated with CIR and RFS was cofounded by high incidences of protocol violations regarding transplants. To address this, we therefore did the sensitivity analyses analyzing only subjects who received their protocol designated therapy. Only *DPEP1* transcript levels remained significantly-associated with CIR and RFS in this cohort. However, we urge caution in accepting this conclusion without external validation.

To understand why *DPEP1* expression might correlate with clinical outcomes we used gain- and loss-of-function cell lines to understand the possible mechanism underlying these associations. *DPEP1* over-expression increased proliferation and survival of B-cell ALL cell lines *in vitro* and in a xeno-transplant model. *DPEP1* knockdown had converse effects.

Dysregulation or aberrant activation of pro-survival and -proliferation signaling cascades in leukemia is usually linked to altered drug sensitivity and resistance^22^. Not surprisingly, we found *DPEP1* over-expression increased resistance to anti-leukaemia drugs.

We also studied changes in several growth- and survival-related pathways in BV173 cells. PI3K-AKT has been identified as a central survival pathway downstream of the B-cell receptor central to normal or neoplastic B-cell development^23^. We detected similar trends in AKT phosphorylation and *DPEP1* expression suggesting an involvement of p-AKT in pro-survival mechanisms mediated by *DPEP1*.

Transcription factors are key regulators of normal and leukaemia cell proliferation, survival and self-renewal^24^. *CREB* is over-expressed and constitutively phosphorylated in several human cancers including AML^25, 26^. We found pCREB levels were increased in parallel with *DPEP1* expression and that down-regulation of *DPEP1* correlated with up-regulation of p53 favouring cell apoptosis rather than proliferation and survival.

There are several important limitations to our study including small sample size, substantial protocol violations and lack of an external validation cohort. Also, subjects were not randomly-assigned to receive tyrosine kinase-inhibitor and/or allotransplant. Because of these limitations, conclusion regards the predictive impact of *DPEP1* of CIR and RFS requires confirmation in a cohort of uniformly-treated subjects.

In summary, our data implicate *DPEP1* expression in the biology of common B-cell ALL in adults. We report clinical correlates, namely an association between high transcript levels and an increased CIR and worse RFS and provide a potential biological basis for these correlations. If confirmed, analyzing *DPEP1* transcript levels at diagnosis could help predict therapy-outcomes. Moreover, regulation of *DPEP1* expression could be a therapy target in B-cell ALL.

## Acknowledgments

This research is supported by grants from the Beijing Municipal Natural Science Foundation [Grant 7192213], the National Natural Science Foundation of China (Grants 81770156, 81570182 and 81270572), the Peking University People’s Hospital Research and Development Funds (Grant RDF2018-02), the Peking University Clinical Scientist Program (Grant BMU2019LCKXJ003) and the Fundamental Research Funds for the Central Universities. RPG acknowledges support from the National Institute of Health Research (NIHR) Biomedical Research Centre funding scheme. Profs. Christian Buske (Ulm Univ.) and Adolpho Ferrando (Columbia Univ.) kindly reviewed the typescript and made helpful suggestions.

Conflict of Interest RPG is a part-time employee of Celgene Corp.

## Author contributions

KYL, GRR designed the project and supervised the typescript preparation. JMZ and YX contributed equally to this work. JMZ and YX performed the experiments, analyzed the data and wrote the manuscript. All the other authors provided the clinical data. All authors have read and approved the final version of this manuscript.

## Abbreviations

ALL: Acute lymphoblastic leukemia
*DPEP1*: Dehydropeptidase1
RT-qPCR: Quantitative real-time polymerase chain reaction
CIR: Cumulative incidence of relapse
RFS: relapse-free survival
MRD: Measurable residual disease
Pro-B-cell ALL: Early precursor B-cell ALL
Pre-B-cell ALL: Precursor B-cell ALL
Com-B-cell ALL: Common B-cell ALL
ROC curve: Receiver-operator characteristic curve

---

**Figure 1S.**
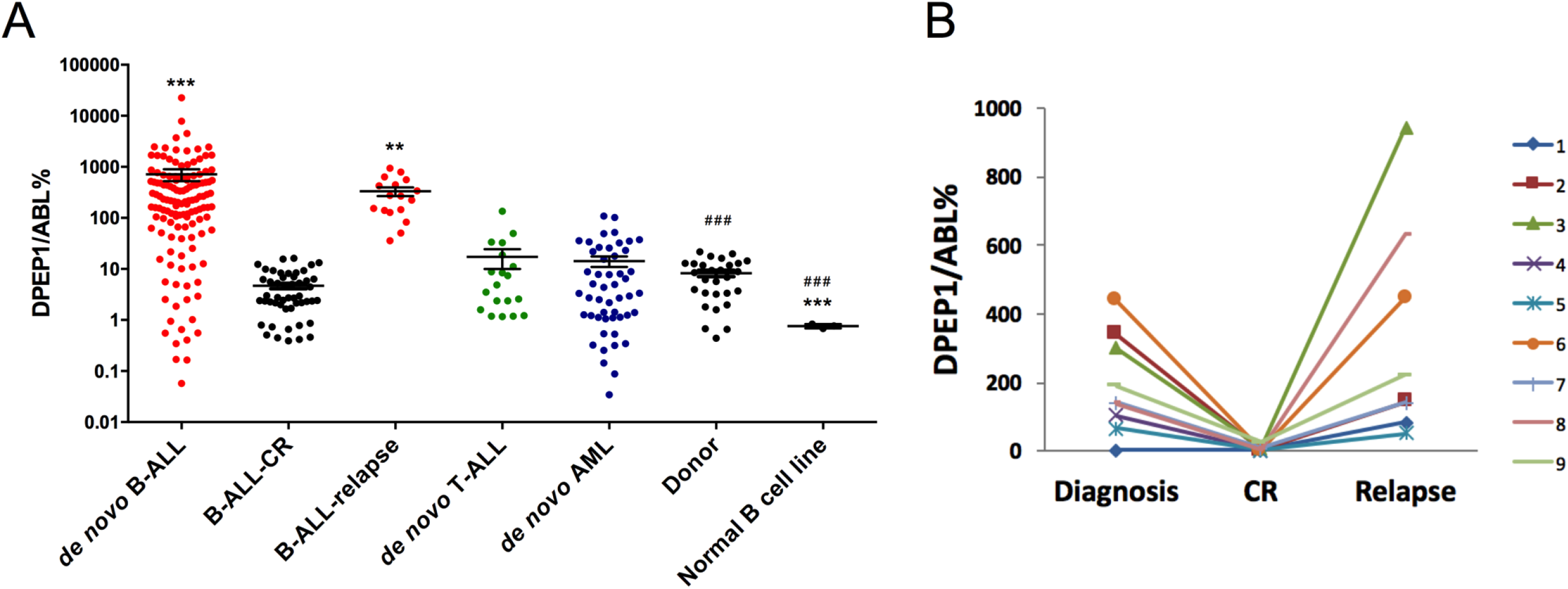
*DPEP1* transcript levels. (A) B-cell ALL at diagnosis, in complete remission, with measurable residual disease (MRD)-test-positive and normals; (B) ROC curve of transcript level of *DPEP1* in normals and subjects with B-cell ALL; (C) *DPEP1* transcript levels in 9 subjects. Error bars, SE. *DPEP1*, dehydropeptidase1; B-cell ALL, B-cell acute lymphoblastic leukemia; CR, complete remission; ROC, receiver-operator characteristic; AUC, area under the curve.

**Table 1S.**
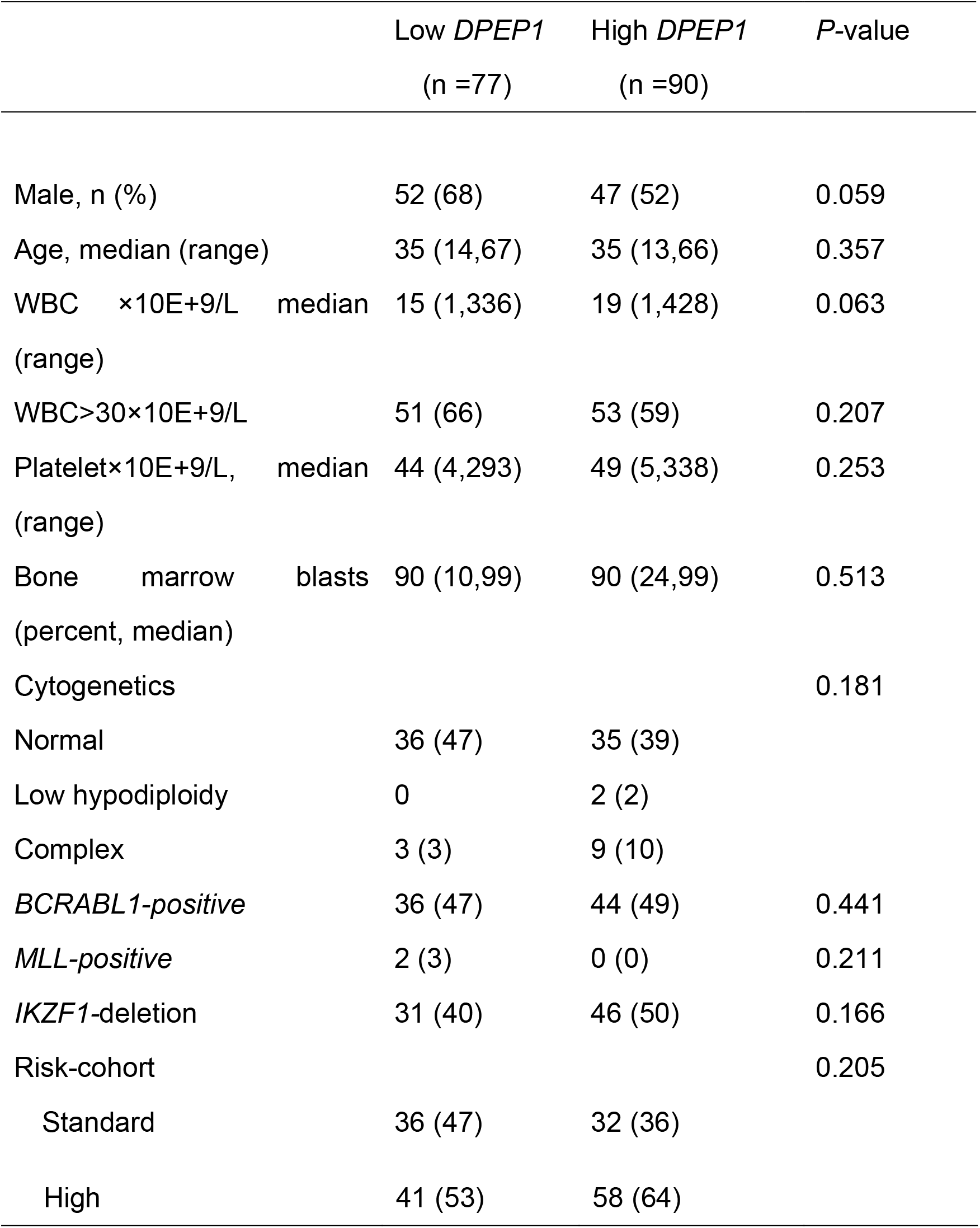
Clinical variables and *DPEP1* transcript levels in subjects with common B-cell ALL.

|  | Low *DPEP1*<br>(n =77) | High *DPEP1*<br>(n =90) | P-value |
| --- | --- | --- | --- |
| Male, n (%) | 52 (68) | 47 (52) | 0.059 |
| Age, median (range) | 35 (14,67) | 35 (13,66) | 0.357 |
| WBC $\times 10E+9/L$ median (range) | 15 (1,336) | 19 (1,428) | 0.063 |
| WBC>30$\times 10E+9/L$ | 51 (66) | 53 (59) | 0.207 |
| Platelet$\times 10E+9/L$, median (range) | 44 (4,293) | 49 (5,338) | 0.253 |
| Bone marrow blasts (percent, median) | 90 (10,99) | 90 (24,99) | 0.513 |
| Cytogenetics |  |  | 0.181 |
| Normal | 36 (47) | 35 (39) |  |
| Low hypodiploidy | 0 | 2 (2) |  |
| Complex | 3 (3) | 9 (10) |  |
| *BCRABL1*-positive | 36 (47) | 44 (49) | 0.441 |
| *MLL*-positive | 2 (3) | 0 (0) | 0.211 |
| *IKZF1*-deletion | 31 (40) | 46 (50) | 0.166 |
| Risk-cohort |  |  | 0.205 |
| Standard | 36 (47) | 32 (36) |  |

|  |  |  |
| --- | --- | --- |
| High | 41 (53) | 58 (64) |

